# Apical expansion of calvarial osteoblasts and suture patency is dependent on graded fibronectin cues

**DOI:** 10.1101/2023.01.16.524278

**Authors:** Xiaotian Feng, Helen Molteni, Megan Gregory, Jennifer Lanza, Nikaya Polsani, Rachel Wyetzner, M. Brent Hawkins, Greg Holmes, Sevan Hopyan, Matthew P. Harris, Radhika P. Atit

**Affiliations:** Department of Biology, Case Western Reserve Univ., Cleveland Ohio, USA; Dept of Genetics, Harvard Medical School, Dept. of Orthopedics, Boston Children’s Hospital, Boston, Massachusetts, USA; Dept. of _Genetics and Genomic Sciences, Icahn School of Medicine at Mount Sinai, New York, New York, USA; Dept. of Developmental Biology, Hospital for Sick Kids, Toronto, Canada

**Keywords:** mouse calvaria, apical expansion, craniosynostosis, Wasl/N-Wasp

## Abstract

The skull roof, or calvaria, is comprised of interlocking plates of bone. Premature suture fusion (craniosynostosis, CS) or persistent fontanelles are common defects in calvarial development. Although some of the genetic causes of these disorders are known, we lack an understanding of the instructions directing the growth and migration of progenitors of these bones, which may affect the suture patency. Here, we identify graded expression of Fibronectin (FN1) protein in the mouse embryonic cranial mesenchyme (CM) that precedes the apical expansion of calvarial osteoblasts. Syndromic forms of CS exhibit dysregulated FN1 expression, and we find FN1 expression is altered in a mouse CS model as well. Conditional deletion of *Fn1* in CM causes diminished frontal bone expansion by altering cell polarity and shape. To address how osteoprogenitors interact with the observed FN1 prepattern, we conditionally ablate *Wasl/N-Wasp* to disrupt F-actin junctions in migrating cells, impacting lamellipodia and cell-matrix interaction. Neural crest-targeted deletion of *Wasl* results in a diminished actin network and reduced expansion of frontal bone primordia similar to conditional *Fn1* mutants. Interestingly, defective calvaria formation in both the *Fn1* and *Wasl* mutants occurs without a significant change in proliferation, survival, or osteogenesis. Finally, we find that CM-restricted *Fn1* deletion leads to premature fusion of coronal sutures. These data support a model of FN1 as a directional substrate for calvarial osteoblast migration that may be a common mechanism underlying many cranial disorders of disparate genetic etiologies.

## Introduction

The roof of the mammalian skull is comprised of a quartet of paired intramembranous bones, the frontal and parietal bones, as well as, in the mouse, the interparietal bone (Carter & Anslow, 2009; Ferguson et al., 2018; Ishii et al., 2015). Where these bones meet, fibrous joints, or sutures, are formed that provide areas for continued growth during postnatal stages (Zhao et al., 2015). Fate mapping in mice shows that the frontal bones originate from the cranial neural crest, while the parietal bones are derived from the paraxial mesoderm (Jiang et al., 2002). However, both the frontal and parietal bone primordia emerge as foci in the supraorbital arch mesenchyme (SOM) above the eye and commit to the osteoblast lineage (Ferguson et al., 2018; Jiang et al., 2002; Yoshida et al., 2008). Through development, the osteoblasts in the calvarial anlagen establish an osteogenic front and expand to the apex of the head to cover the brain (Fan et al., 2016). The mechanisms of how the calvarial osteogenic progenitors expand to their respective locations in a determinative manner and consistently form without deformation are not understood.

*Ex-utero* DiI-labeling as well as inducible genetic lineage mapping of the mouse SOM cells between embryonic day (E)10.5-13.5, showed that SOM cells contribute to bone tissue over the frontal and parietal bones and adjacent dermal mesenchyme by E16.5 (Deckelbaum et al., 2006; Tran et al., 2010; Yoshida et al., 2008). The distribution pattern of labelled cells suggest that active cell movement may play a role in expansion of calvaria. Iseki et al. (1999) and Opperman et al. (2000) demonstrated that the population of osteoprogenitors located at the osteogenic fronts of the bone anlagen are highly proliferative. However, Lana-Elola et al. (2007) found that inhibiting cell proliferation of the calvarial explants *ex-vivo* diminished parietal bone growth apically by less than 20%. These data suggest that proliferation is not the prominent driver of apical expansion of the calvaria. End addition of small population of ephrin-Eph positive ectomesenchyme cells directly at the osteogenic front are complementary mechanisms for growth, but they are not sufficient to drive apical expansion (Ting et al., 2009). We hypothesize that the apical expansion of osteoprogenitors over a long-distance results from directional cell migration.

Directional migration of calvarial osteoblast progenitors towards the apex would require guidance cues to establish coordinated cell activity, perhaps graded across the embryonic cranial mesenchyme (CM). Using RNA-seq analysis at E12.5, Dasgupta et al. (2019) found that genes related to cell adhesion and extracellular matrix (ECM)-receptor interaction pathways are differentially expressed in the in previously established mesenchyme population located apically to the SOM. The interstitial ECM is a set of macromolecules such as collagen, fibronectin and laminin, and polysaccharide components. During embryonic tissue development, the interstitial ECM plays dual roles in providing either structural support and/or regulation of signaling pathways important for cell movement and tissue growth (Walma & Yamada, 2020). Differential expression of ECM-receptor interactions suggests potential directional migration cues and substrate to guide apical expansion of calvarial osteoblast progenitors. Even with these insights, there remains no clear evidence for active migration or which tissue signals promote directional cell movement over a long distance from above the eye to the apex of the head. A fundamental understanding of early calvarial morphogenic processes could provide new insights into the etiology of dysplasia and congenital defects of the skull such as craniosynostosis.

In this study, we assessed whether Fibronectin (FN1), a key interstitial ECM protein that was identified as having a role in directional cell movement *in vitro* (Lo et al., 2000; Shellard & Mayor, 2020), acts as a substrate for calvarial osteoblasts in both apical expansion of calvaria as well as the maintenance of the sutures in embryonic mice. We demonstrate that FN1 protein serves as a substrate and is required for specific cellular behaviors of calvarial osteoblasts during apical expansion. We find that deletion of FN1 in calvarial mesenchyme results in premature fusion of the coronal suture at high penetrance. F-actin can serve as anchor points that mediate ECM interactions. Targeted deletion of *Wasl*, an F-actin nucleating factor (Caswell & Zech, 2018), in cranial mesenchyme *in vivo* leads to the specific failure of calvarial osteoblasts to migrate. These results are consistent with lamellipodia-mediated cell migration, potentially across a graded FN1 substrate, leading to proper patterning of the skull roof, and provide a new mechanistic model of calvarial morphogenesis.

## Results

### FN1 expression is graded along the basal-apical axis in cranial mesenchyme

To assess potential asymmetric cues present in the developing skull roof, we examined the expression of interstitial ECM proteins during early stages of SOM formation (E12.5-13.5) that might serve as substrates for cell movement (Walma & Yamada, 2020). We initially visualized pan-collagen unfolded and denatured chains by staining with fluorescent Collagen Hybridizing Peptide on E12.5 coronal sections (**Figure 1A**). Detection of these products was not regionally different in mesenchymal populations (**Figure 1A**). Consistent with this result, Collagen III was not highly expressed in CM, but it was visible in the meninges (**Figure 1 B**). Additionally, using a broad assessment of mineralized collagen pattern by RGB and Masson’s Trichrome histochemical stains, showed that mineralized collagen is also not prominent in the CM at E13.5 (**Figure 1D, Supplementary Figure S1C)**. Laminin, which is a dominant ECM protein in basement membranes and a component of the fibrous networks of ECMs, was expressed in the developing meninges and blood vessels in the CM at E12.5. However, laminin was not visible in the CM (**Figure 1C**). Similarly, the expression of other fibrous ECM molecules such as sulfated proteoglycans and Elastin at these stages did not show enrichment (**Supplementary Figure S1B)**. In contrast, we found that Fibronectin1 (FN1) protein was highly enriched in the CM. Notably, its expression was graded towards the apex suggesting that it may provide external directional cues for calvarial osteoblasts **(Figure 1E-1G & 2F**). We independently validated the graded expression of FN1 in cranial mesenchyme by assessing the expression of Fibulin 1 protein (**Supplemental Figure S1A**), which is associated with assembled FN1 protein fibers in the extracellular matrix (Balbona et al., 1992). We found that Fibulin1 protein was also enriched in the CM toward the apex, similar to the graded expression of FN1. Together, these results show that among the expression of the core fibrous interstitial ECM components (Walma & Yamada, 2020) the expression of FN1 stands out because it is enriched and graded towards the apex in CM, in the direction of calvarial expansion (**Figure 1H**).

**Figure 1:**
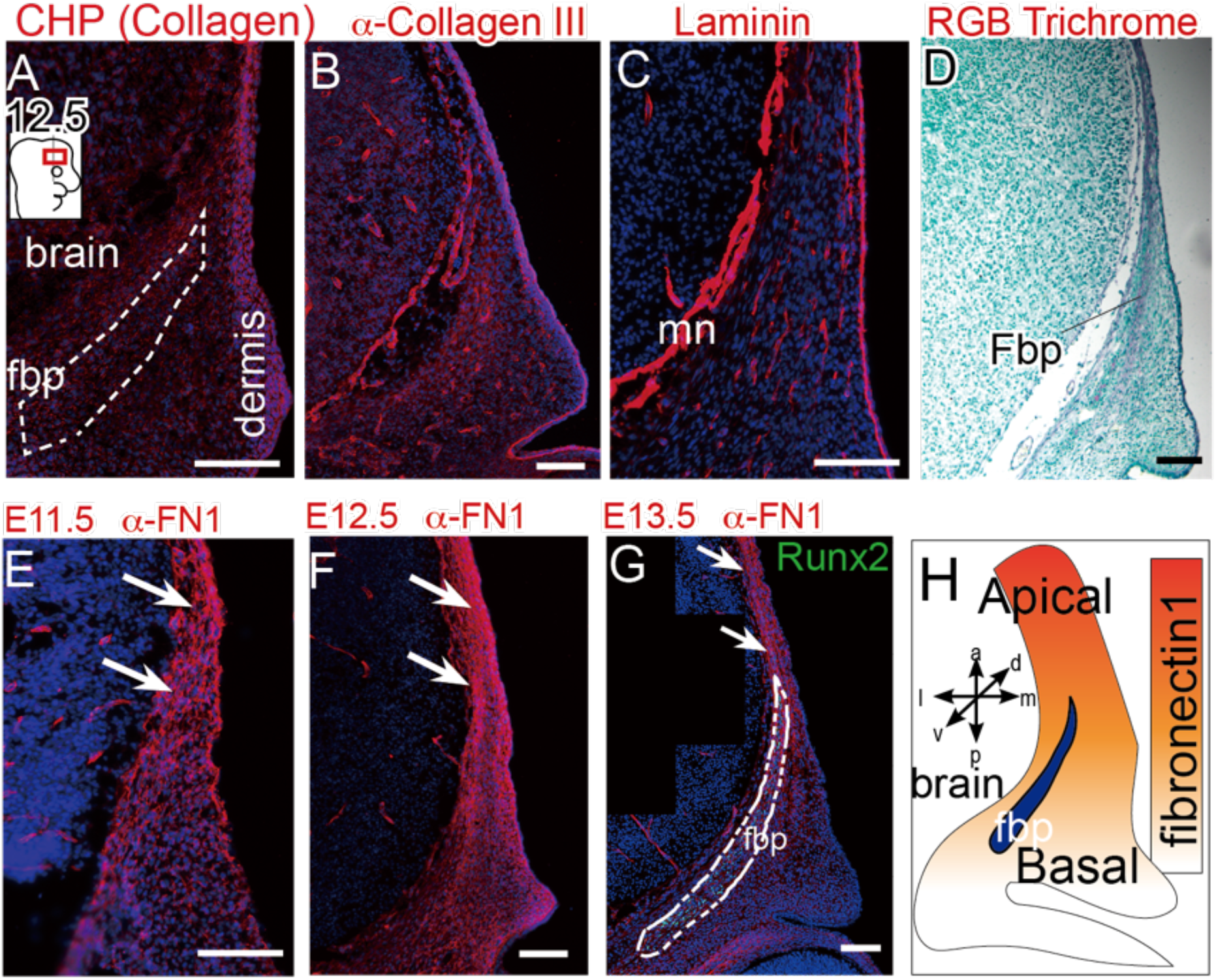
Fn expression is graded towards the apex in cranial mesenchyme, in contrast to other core ECM proteins. (A) Immunofluorescence of unfolded collagen chains in E12.5 coronal sections. Immunofluorescence of collagen III (B), Laminin (C), RGB Trichrome staining for collagen (D) in E12.5 coronal sections. (E, F, G) Immunofluorescence of FN1 in coronal sections showing graded FN1 protein expression in cranial mesenchyme at E11.5-13.5. N≥3 controls; 3 mutants. (H) Schematic depicting the graded FN1 substrate towards the apex of the cranial mesenchyme. Nuclei were labeled with DAPI in blue. CHP, collagen hybridizing peptide; FN1, Fibronectin 1; fbp, frontal bone primordia. mn, meninges. Scale Bar= 100 μM.

### Elevated FN1 levels accelerate apical expansion of calvarial osteoblasts in developing frontal bone primordia

The graded expression of FN1 in the cranial mesenchyme along the baso-apical axis could function as a tissue level directional substrate for guiding the calvarial osteoprogenitors to their destinations. Humans with Apert syndrome have calvarial defects and CS, and suture fibroblasts from these patients secrete higher FN1 in culture (Bodo et al., 1997). We visualized FN1 protein expression in CM of the Apert syndrome mouse model which recapitulates human calvarial defects (L. Chen et al., 2003; Holmes & Basilico, 2012; Wang et al., 2005). Expression along the baso-apical axis in the frontal bone primordia at E13.0 showed significantly elevated FN1 expression in all three regions of the CM in Apert mice compared to controls (**Figure 2A, B & E**). However, FN1 expression in Apert mutants remained graded towards the apex (**Figure 2E**). To further investigate the relation between FN1 expression in the CM with the apical expansion of calvarial bones, we quantified the apical extension of alkaline phosphatase, an early osteogenic marker, in frontal bone primordia in E13.0 (Ishii et al., 2015). Relative to litter-matched control embryos, we found the relative length of frontal bone primordia in Apert mutants, and thus apical expansion of calvarial osteoprogenitors, is significantly increased (**Figure 2C, D & F**). Together, these data suggest that elevated expression of FN1 in the CM correlate with increased expansion of the embryonic frontal bone primordia.

**Figure 2.**
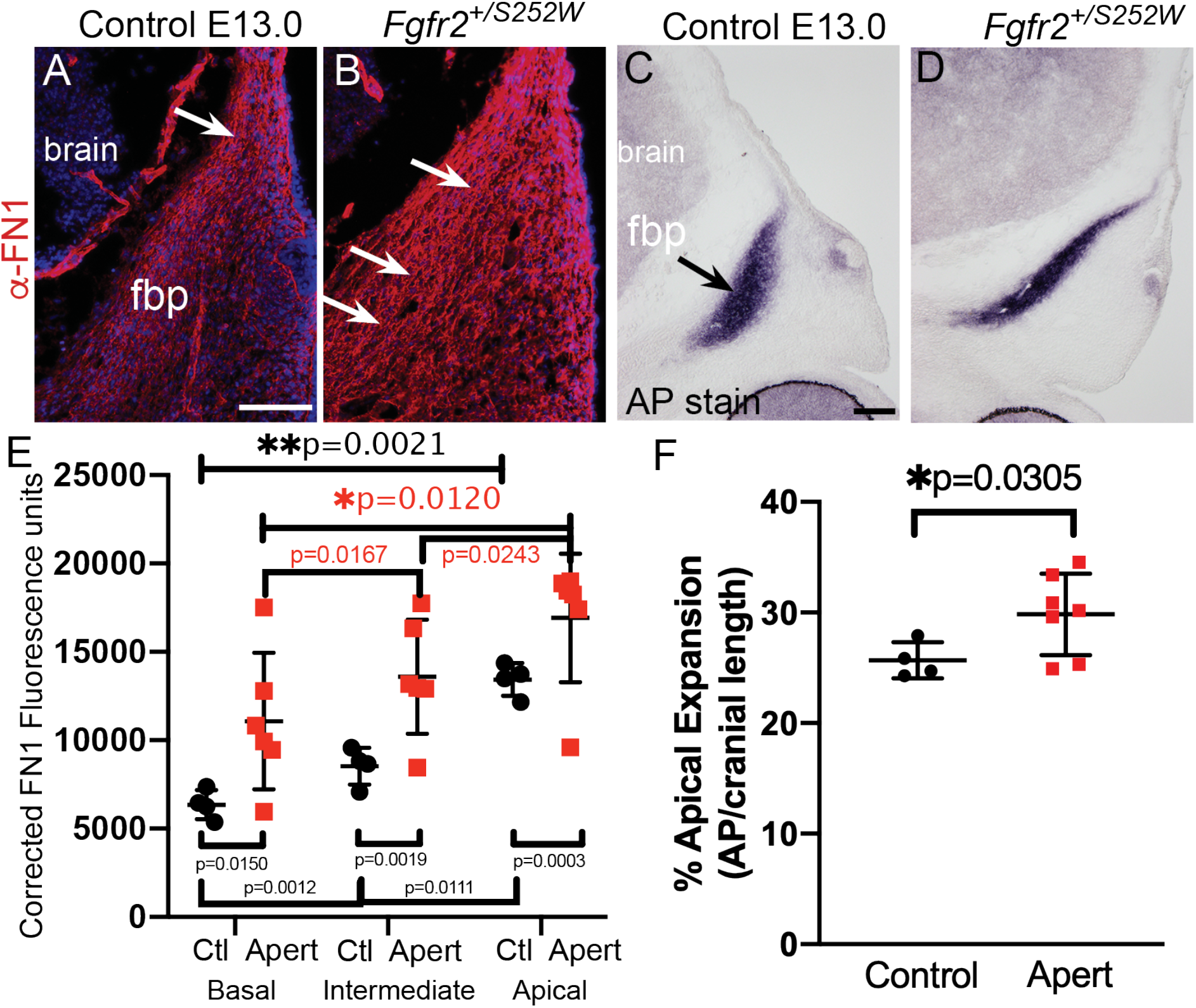
FN1 is elevated in CM towards the apex in the Apert syndrome mouse model. (A, B, E) Immunofluorescence of FN1 in E13.0 coronal sections showing elevated FN1 in Apert mutants relative to controls. Quantification of FN1 expression in cranial mesenchyme shows the graded FN1 expression in both controls and Apert mutants. The FN1 expression level is increased in Apert mice relative to controls. N=4 controls; 6 mutants (C, D, F) Alkaline phosphatase staining in E13.0 coronal sections showing increased apical length of frontal bone primordia relative to controls. N=4 controls; 7 mutants. Fbp, frontal bone primordia; AP, alkaline phosphatase; Ctl, control. Scale Bar= 100 μM.

### FN1 matrix is required for apical expansion of calvarial progenitors

Fibronectin1 plays many crucial roles in tissue growth and morphogenesis during development and FN1-Integrin interactions are required for differentiation of osteoblasts *in vitro* (Globus et al., 1995; Yamada et al., 2019). Whether and how FN1 functions in the cellular behavior and differentiation of osteoblasts during calvarial development and patterning *in vivo* is unknown. To address the role FN1 in calvarial development, we conditionally deleted *Fn1* in the CM in mouse between E10-11.0 using tamoxifen-inducible *Pdgfrα-CreER* mediated recombination (CM-*Fn*^*fl/fl*^) (Ferguson et al., 2018) (**Figure 3A**). We validated the efficiency of *Fn1* deletion in the cranial mesenchyme at E13.5 and found a significant decrease of *Fn1* mRNA by qPCR and loss of FN1 protein expression by immunofluorescence staining in CM (**Figure 3B, C & D**). We examined the cell proliferation index of calvarial osteoblasts between E12.5-E14.5 in the frontal bone primordia and found it was not significantly different between controls and CM-*Fn*^*fl/fl*^ mutants (**Figure 3E**). Cell survival in the cranial mesenchyme at E12.5 was also not altered in CM-*Fn*^*fl/fl*^ embryos (**Supplementary Figure S3D, E**). Next, we tested if osteoblast lineage specification and commitment was affected by *Fn1* deletion in the CM (Nakashima et al., 2002). The percentage of RUNX2^+^ osteoblast progenitors at E12.5 in three non-overlapping regions in the frontal bone primordia was comparable in the control and CM-*Fn*^*fl/fl*^ embryos (**Figure 3I, F**). At E14.5, the percent of OSX^+^ committed osteoblasts were comparable in control and CM-*Fn*^*fl/fl*^ mutants along the entire baso-apical axis of the frontal bone primordia (**Figure 3I, J**).

**Figure 3.**
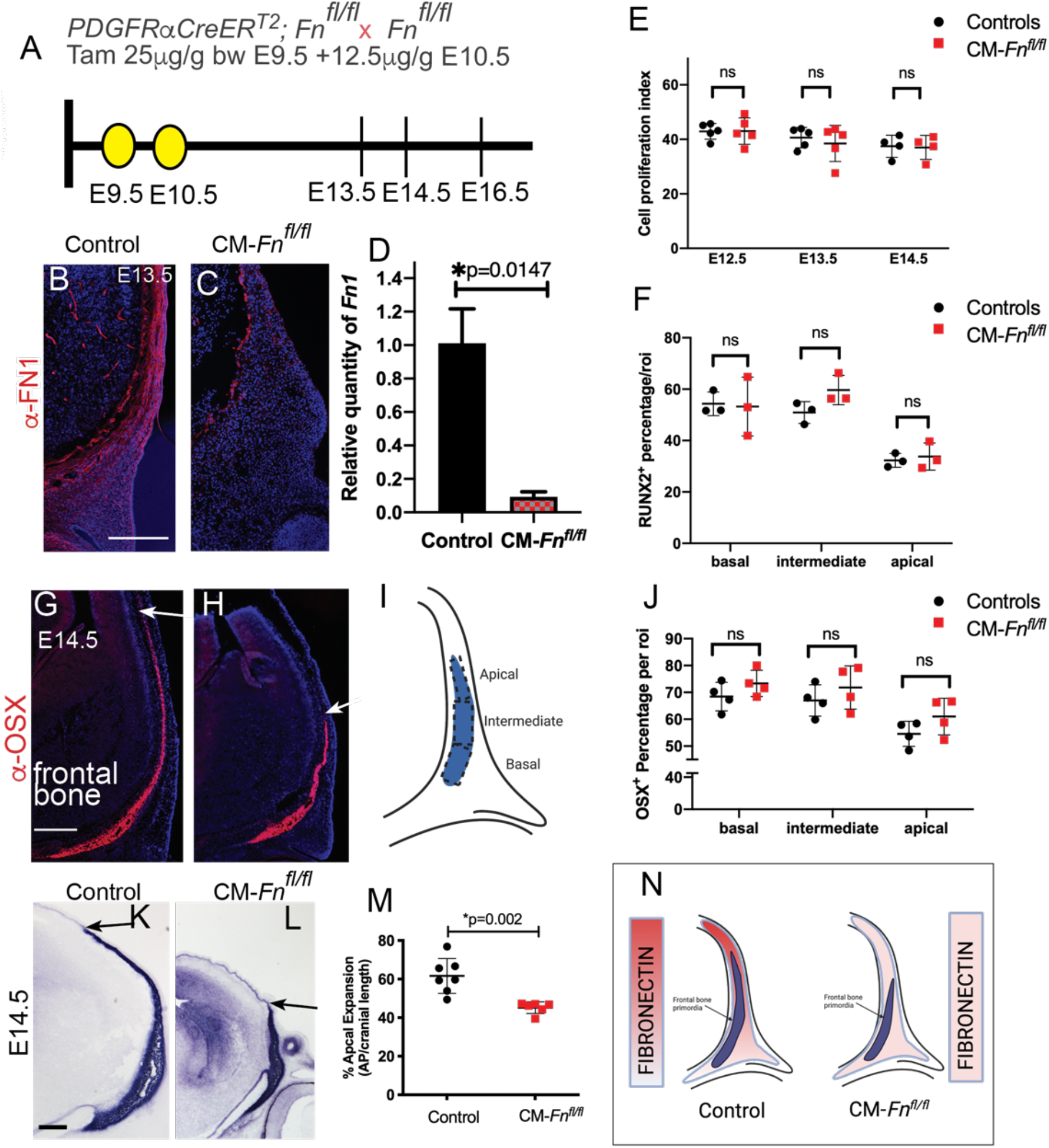
Loss of FN1 results in decreased apical expansion of frontal bone primordia at E14.5, without perturbing cell proliferation. (A). Schematic of tamoxifen inducible deletion of FN1 in the SOM at E9.5 and E10.5 and the schematic of the region of interest in coronal section. (B, C) Immunofluorescence of FN in E13.5 coronal sections and (D) RT-qPCR analysis showing diminished FN expression and relative quantity of mRNA transcripts for Fn in E13.5 CM-*Fn*^*fl/fl*^ mutants relative to controls demonstrating deletion efficiency of gavage regimen. (E) Proliferation index of calvarial osteoblasts among controls and CM-*Fn*^*fl/fl*^ mutants at E12.5-E14.5 is comparable. (F) Percentage of calvarial osteoprogenitor marker RUNX2^+^ cell in the region of interest in different regions of cranial mesenchyme shows no significant difference between controls and CM-*Fn*^*fl/fl*^ mutants at E12.5. (I) Schematic depicting the measurements of RUNX2^+^ and OSX^+^ cell percentages in apical, intermediate, and basal regions of the frontal bone primordia. (G, H) Immunofluorescence of calvarial osteoblast marker OSX showing decreased apical expansion of frontal bone in E14.5 CM-*Fn*^*fl/fl*^ mutants relative to controls. (J) Percentage of OSX^+^ cell in the region of interest in different regions of cranial mesenchyme shows no significant difference between controls and CM-*Fn*^*fl/fl*^ mutants at E14.5. (K, L, M) The apical length of the frontal bone primordia is decreased in CM-*Fn*^*fl/fl*^ mutants relative to controls at E14.5. (N) Summary schematics showing that the apical length of frontal bone primordia is decreased in FN1 deleted cranial mesenchyme. N=3-7 controls; 3-5 mutants. Scale Bar = 100 μM.

We observed that the spatial domain of OSX expression was markedly diminished in the absence of FN1 (**Figure 3G, H**). Next, we measured the apical extension of the frontal bone primordia marked by alkaline phosphatase activity (ALP) at E14.5 (**Figure 3K, L, & M**) and at E13.5 (**Supplementary Figure S3A, B, & C**). We found that there was a significant decrease in the apical extension of the frontal bone primordia in CM-*Fn*^*fl/fl*^ mutants relative to stage matched controls starting at E14.5 but not at E13.5 (**Figure 3K-M**). Consistently, cell density of the frontal bone anlagen was significantly increased in CM-*Fn*^*fl/fl*^ mutants relative to the controls, notably at the basal region of the frontal bone primordia (**Supplementary Figure S3F**). All together, these results suggest that lack of FN1 expression in CM reduces apical expansion of the frontal bone progenitors, without perturbing cell proliferation, osteogenesis, or cell survival (**Figure 3N**).

### FN1 is required for polarization and elongation of cranial mesenchyme cells along graded aspect of developing skull

In order to drive tissue morphogenesis and extension, cells become polarized and change shape, which can be coordinated by cues from the ECM (Yamada et al., 2019). However, it is not known if calvarial progenitors are polarized during apical expansion or if FN1 is required for cellular polarity. The asymmetric position of the condensed Golgi complex is a well-known indicator of cellular polarity and is a required aspect of directed cell movement in a variety of cells in culture (Drabek et al., 2006; Yadav et al., 2009). We measured the angle between the Golgi and the nucleus in E13.5 calvarial osteoblasts along the baso-apical axis in the frontal bone primordia (**Figure 4A-C**). The Golgi angle distributions of the control frontal bone primordia have a bi-directional bias in distribution (**Figure 4D**). However, in the CM-*Fn1*^*fl/fl*^ mutants, the distribution of the Golgi and nucleus angle of calvarial osteoblasts was directed rearward toward the eye and significantly different in both the basal and intermediate regions of frontal bone primordia compared to controls (**Figure 4D**). In the apical region of the frontal bone primordia, the Golgi angle distribution was highly biased to the apex at 90 degrees in controls and CM-*Fn*^*fl/fl*^ mutants (**Figure 4D**). This is consistent with earlier studies that show that migrating cell types often position Golgi rearward of the nucleus and this positioning is greatly impacted by geometric constraints in fibronectin substrate (Pouthas et al., 2008; Serrador et al., 1999). Given that mesenchyme cells have uneven boundaries and nuclei shape tracks cells morphology with high fidelity, we examined nuclei shape morphology of calvarial osteoblasts (Chen et al., 2015). In E13.5 controls, calvarial osteoblasts nuclei become progressively elongated along the baso-apical axis in the controls. The nuclei length-width aspect ratio of calvarial osteoblasts in the basal region was between 1:1-4:1 and between 1:1-7:1 in the apical regions (**Figure 4J**). In CM-*Fn*^*fl/fl*^, the distribution of length-width aspect ratio was significantly less in basal and apical regions (**Figure 4J**). The calvarial osteoblasts in CM-*Fn*^*fl/fl*^ mutants had decreased length-width ratio indicating the cells are less elongated throughout the frontal bone primordia. Collectively, these findings indicate that apical expansion of the frontal bone at the tissue level occurs by cellular polarization in calvarial osteoblast that are most elongated and these cellular behaviors in the basal region is dependent on FN1 expression in the CM.

**Figure 4.**
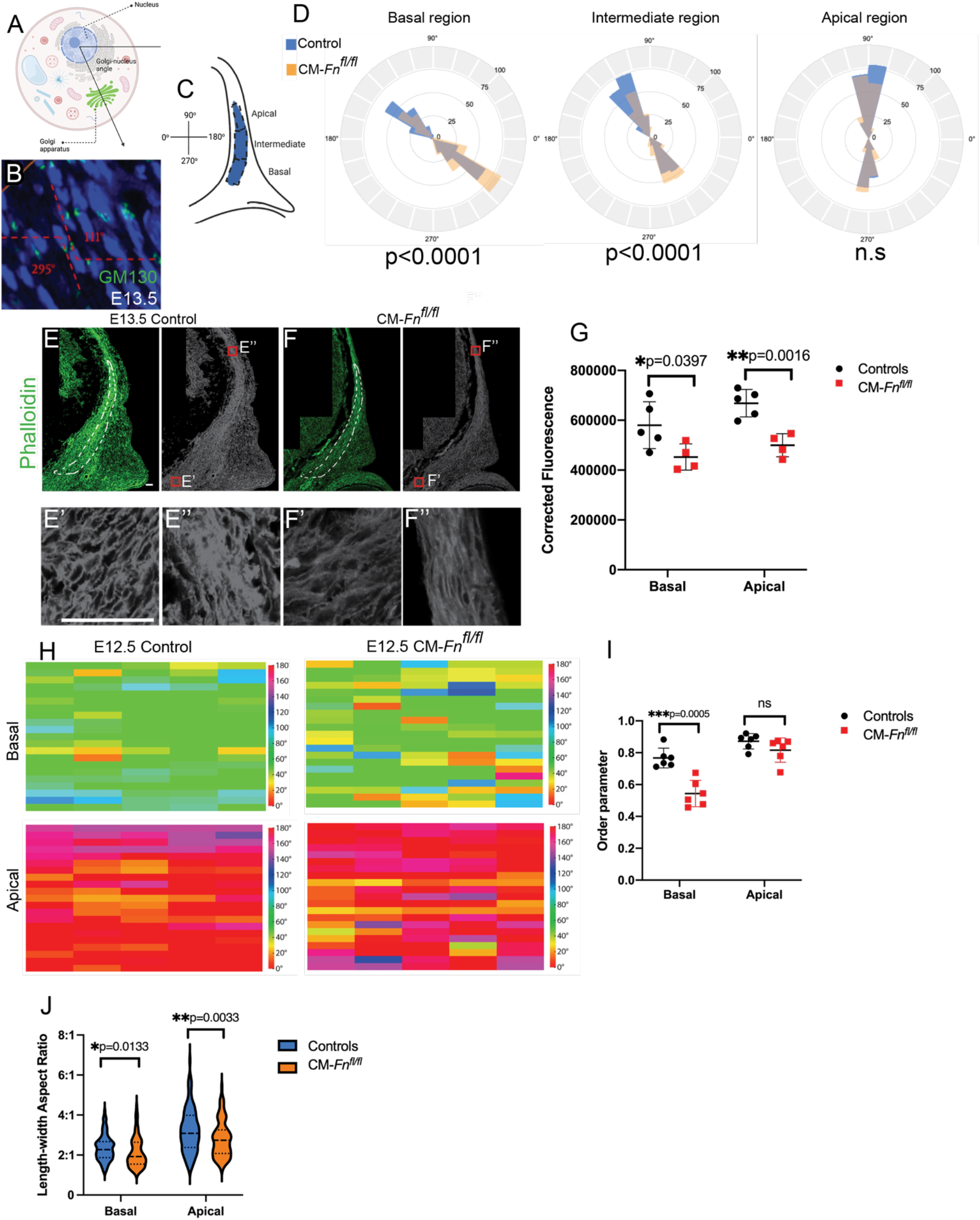
FN1 deletion disturbs the cellular polarity, cell shape, and actin distribution of calvarial osteoblasts. (A, B, C) Schematics depicting the area of interest in coronal sections and the analysis of cellular polarity through GM130 immunofluorescence. (D) Windrose plots showing the cellular polarity distribution of calvarial osteoblasts in basal and intermediate regions of cranial mesenchyme is perturbed in CM-*Fn*^*fl/fl*^ mutants relative to controls at E13.5. N = 4 controls, 4 mutants. (E, E’, E’’, F, F’, F’’, G) The staining of phalloidin showing diminished level of F-actin in CM-*Fn*^*fl/fl*^ mutants at E13.5 relative to the controls. N = 5 controls, 4 mutants. Dash lines label the region of frontal bone primordia. (H, I) Heat map and Order parameter representing vector angles of the fibrillar alignment of F-actin shows anisotropy is significantly less in CM-*Fn*^*fl/fl*^ mutants in the basal region and comparable to controls in the apical region at E12.5. The heat maps were generated by the data collectively obtained from a fixed sized-ROI in apical and basal regions of the frontal bone primordia of three biological replicates. For the order parameters, data were obtained from the apical and basal regions of the frontal bone primordia in two histological sections of three biological replicates, respectively. (J) The length-width ratio of calvarial osteoblasts of CM-*Fn*^*fl/fl*^ mutants is significantly lower in basal and apical regions relative to the controls at E13.5. N =4 controls, 4 mutants. Scale Bar = 50 μM.

### Fibronectin is required for enrichment of F-actin in the cranial mesenchyme

Actin filaments are part of the force generating complex used for cell migration (Walma & Yamada, 2020; Yamada & Sixt, 2019). During cell movement, mesenchymal cells can have enriched and aligned contractile stress fibers of actin cytoskeleton associating with the ECM through polarized actin-rich focal adhesions to promote movement (Walma & Yamada, 2020; Yamada & Sixt, 2019). To determine if actin cytoskeleton expression and organization is dependent on extracellular FN1 expression, we visualized and analyzed the topography of filamentous-actin (F-actin) in the CM by phalloidin staining. In E12.5 and E13.5 controls, F-actin expression was highly enriched in the CM in the domain of the frontal bone primordia (**data not shown, Figure 4E, F)**. In individual mesenchymal cells in the basal region of the frontal bone primordia, F-actin appears to be distributed at the cell boundary (**Figure 4E’, F’**). In the apical region, F-actin appears to be distributed throughout the cell (**Figure 4E’’, F’’**). Relative fluorescent intensity measurement of phalloidin in regions of interest shows a significant decrease in basal and apical regions of the frontal bone primordia in CM-*Fn1*^*fl/fl*^ mutants (**Figure 4E, F, G**). Next, we generated a vector map of actin fibers in regions of interests with Alignment by Fourier Transform (AFT) algorithm (Marcotti et al., 2021). The heat map of a representative individual and compiled ROIs and the median order parameter representing the vector angles of F-actin fibers shows the basal and apical region of the controls are anisotropic and highly aligned to each other (**Figure 4H, I, Supplementary Fig. S4A, B**). The median order parameter approaches 1 in the apical region to signify perfect alignment (**Figure 4I**). The order parameter in the basal region is significantly lower in the CM-*Fn1*^*fl/fl*^ mutants than control, showing, the alignment is significantly less in the basal region, indicating a decrease in anisotropy in F-actin fiber organization. Thus, F-actin distribution and the anisotropy in alignment in calvarial osteoblasts suggest that actin-dependent cell contractility can promote the apical movement of the calvarial osteoblasts, and by extension, frontal bone formation; the process are dependent on presence of graded extracellular FN1.

### The apical expansion of the frontal bone is diminished in *Wnt1Cre2; Wasl*^*fl/fl*^ mutants

To directly test if lamellipodia-dependent cell migration is a mechanism underlying apical expansion of calvarial osteoblasts *in vivo*, we conditionally deleted WASP-Like Actin Nucleation Promoting Factor (*Wasl*), or N-Wasp, a gene encoding a members of Wiskott-Aldrich syndrome protein family, a known effector of CDC42 and ARP2/3 which are key regulators of actin nucleation in lamellipodia (Hüfner et al., 2002; Watson et al., 2017). ARP2/3 is specifically required to direct differential dynamics of lamellipodia for cells moving up along a fibronectin gradient *in vitro* by haptotaxis (King et al., 2016). Conditional deletion of *Wasl* by *Wnt1Cre2* line specifically removes *Wasl* function in mouse cranial neural crest cells as they emerge from the neural tube prior to forming progenitors of the frontal bone and other facial structures. We found that conditional deletion of *Wasl* lead to distinct craniofacial phenotypes including cleft palate and calvarial defects. MicroCT imaging of *Wnt1Cre2; Wasl*^*fl/fl*^ mutants at P0 show mineralized but dramatically reduced size of frontal bone (**Figure 5A**), while parietal bones were generally unaffected, consistent with different embryonic origins of these bones. The leading edge of the *Wasl* mutant frontal bones lacked the irregular border of mineralized bone spicules seen in controls. Consistent with our data from conditional deletion of FN1 substrate, calvarial osteoblast lineage commitment indicated by OSX expression was visible in frontal bone primordia, but the apical expansion of the frontal bone primordia was markedly diminished in *Wasl* mutants compared to controls at E14.5 (**Figure 5B, C**). The overall cell proliferation index of the osteoblasts was comparable at E14.5, number of cells in a fixed field was significantly higher in the frontal bone primordia of *Wnt1Cre2; Wasl*^*fl/fl*^ embryos was increased (**Supplementary Figure S5**). At earlier embryonic stages (E12.5), we did find that the cell proliferation index of RUNX2^+^ calvarial osteoblast progenitors was elevated in the basal region of the frontal bone primordia in *Wnt1Cre2;Wasl*^*fl/fl*^ mutants compared to stage matched controls. However, in the apical regions cell proliferation remained comparable as seen at later stages (**Figure 5D, E & F)**. Further, cell death at E12.5, assayed by expression of cleaved caspase3 expression, was not apparent in cranial mesenchyme of controls or mutants (**data not shown)**. These data suggest that a primary factor leading to the mutant phenotype is failure of calvarial osteoblasts to expand apically.

**Figure 5.**
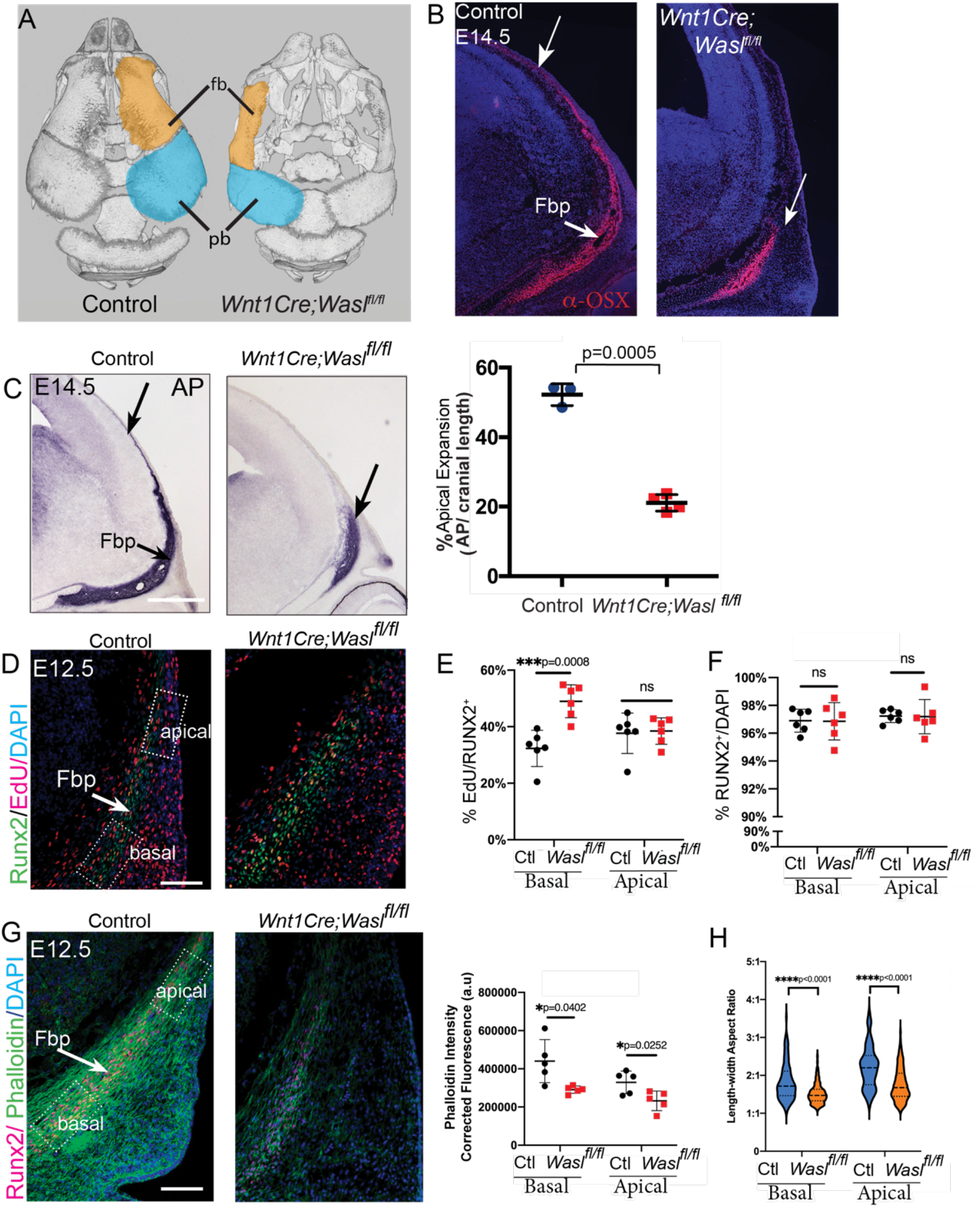
The apical expansion and actin distribution of frontal bone primordia is diminished in *Wnt1Cre; Wasl*^*fl/fl*^ mutants. (A) Dorsal view of μ-CT of calvaria at E17.5, demonstrating diminished apical expansion of the frontal bones (orange) in the mutants. N = 4 controls; 4 mutants. (B and C) Coronal sections in the frontal bone with indirect Immunofluorescence for OSX (B), and alkaline phosphatase staining (C) at E14.5. Quantification showing that the apical length of frontal bone primordia is decreased in *Wnt1Cre; Wasl*^*fl/fl*^ mutants relative to the controls. N = 3 controls; 4 mutants. (D) Immunofluorescence for cell proliferation with EdU, osteoprogenitor marker RUNX2 and counterstained with DAPI at E12.5. Quantification showing increased cell proliferation index in the basal region of mutants relative to the controls at E12.5 (E), with intact cell specification of the osteoprogenitor (F). N= 6 controls; 6 mutants. (G) Coronal sections in the frontal bone primordia stained for F-actin with Phalloidin and immunofluorescence with RUNX2. Corrected fluorescence analysis shows diminished intensity of phalloidin intensity in *Wnt1Cre; Wasl*^*fl/fl*^ mutants relative to the controls. (H) Quantification for length-width aspect ratio showing that the osteoblasts in the *Wnt1Cre; Wasl*^*fl/fl*^ mutants are less elongated. N = 4 controls; 4 mutants. All controls and mutants are shown at the same scale. Scale bar = 100μM. Fb, frontal bone; Fbp, frontal bone primordia; pb, parietal bone.

To assess the impact of *Wasl* deletion on cell behavior, we visualized F-actin distribution and nuclei shape of calvarial osteoblasts in E12.5 frontal bone primordia. Compared to controls, F-actin expression was markedly diminished in the RUNX2^+^ calvarial osteoblasts and the adjacent *Wnt1Cre2* derived mesenchyme in conditional *Wasl* mutant embryos (**Figure 5G**). Consistently, the length-width ratio of nuclei in the basal and apical regions of the frontal bone primordia were significantly decreased in *Wasl* mutants compared to controls (**Figure 5H**). Thus, similar to *Fn1* conditional mutants, *Wasl1-*dependent, actin nucleation in lamellipodia is required for apical expansion of the frontal bone primordia without perturbing overall osteogenesis, survival, and proliferation. Collectively, these analyses expose a new role for FN1 matrix substrate and *Wasl* dependent-lamellipodia in calvarial osteoblasts during apical expansion of the skull roof.

### Deficiencies in FN1 lead to coronal craniosynostosis

Calvarial bone fibroblasts from young children with CS have dysregulated FN1 expression *in vitro* (Baroni et al., 2002; Bodo et al., 1997). This raises the question of whether maintenance of suture patency is associated with FN1 expression in addition to calvarial osteoblast migration. We investigated whether FN1 substrate is also required for the maintenance of sutures. In control E16.5 mouse embryos the coronal suture was patent (**Figure 6A**). In contrast, in CM-*Fn*^*fl/fl*^ mutants that survive till E16.5, we observed alizarin red staining in the coronal suture space showing premature fusion unilaterally as well as bilaterally (**Figure 6B**). The penetrance of the craniosynostosis phenotype in FN1-deficient CM cells was approximately 80% (**Figure 6C)**. In transverse sections of CM-*Fn*^*fl/fl*^ mutants, we confirmed ectopic mineralization in the coronal suture space showing inappropriate fusion of the sutures (**Figure 6D-F**). Relative to the stage matched controls, we found the frontal bone area is decreased and anterior metopic (frontal) fontanelle distance between the paired frontal bones was significantly increased in CM-*Fn*^*fl/fl*^ mutants, suggesting these phenotypes are linked (**Figure 6G-L**). Together, these findings indicate that FN1 function in the cranial mesenchyme is required to preserve coronal suture patency while promoting apical expansion of the frontal bone primordia.

**Figure 6.**
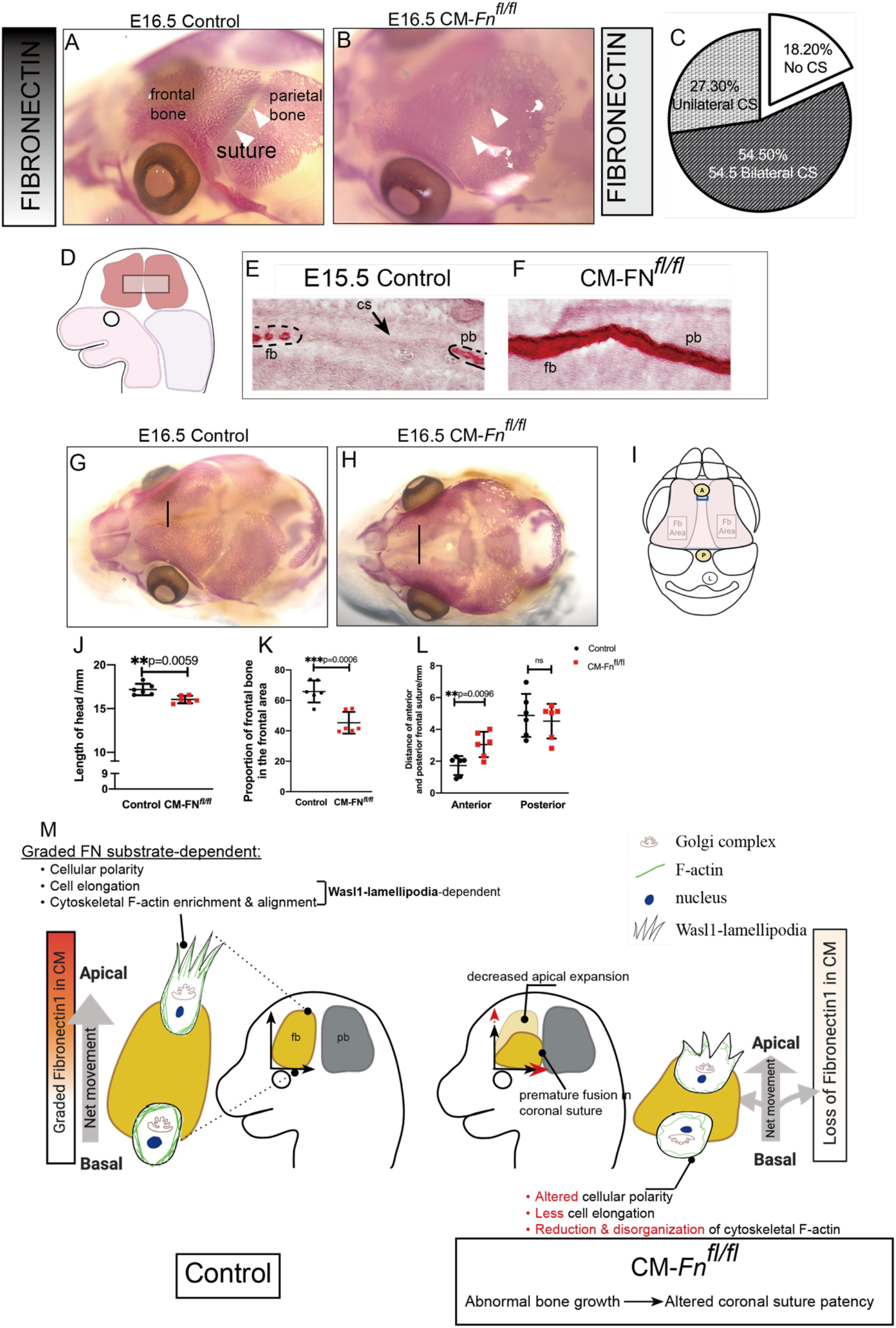
Loss of FN1 in cranial mesenchyme causes craniosynostosis in coronal suture and affects the growth of the frontal bone. (A, B, C) Alizarin skeletal staining of E16.5 controls and CM-*Fn*^*fl/fl*^ mutants showing craniosynostosis in coronal suture at approximately 80% penetrance. (D) Schematic of the coronal suture depicting the area of interest for the transverse sections. (E, F) Alizarin staining of E15.5 transverse sections showing the ectopic mineralization in the coronal suture space in CM-*Fn*^*fl/fl*^ mutants. (G, H) Alizarin skeletal staining showing the increased distance between the frontal bones in CM-*Fn*^*fl/fl*^ mutants at E16.5. (I) Schematic depicting the histomorphometry analysis for the skeletal staining. (J, K, L) Quantifications of the skeletal staining showing the width between anterior frontal bones is increased whereas the proportion of frontal bone to the frontal area (labeled in pink) is decreased in CM-*Fn*^*fl/fl*^ mutants which have shorter skull length. N=6 controls; 6 mutants. (M) A schematic of working model illustrating the role of FN1 in apical expansion of frontal bone primordia and maintenance of coronal suture patency. Fb, frontal bone; pb, parietal bone; cs, coronal suture. For images of wholemounts, A and B, C and D were obtained at the same magnification, respectively.

FN1 is required for efficient expansion of calvarial osteoblast expansion and to maintain suture patency. Calvarial osteoblast behaviors such as polarization, elongated cell shape, and actin cytoskeleton enrichment demonstrate that they use FN1 as a substrate for apical expansion. Thus, the overlapping phenotypes of *Fn1* and *Wasl1* mutants suggest that they are working together in a biological process for apical expansion allowing us to build a new model for formation of the skull roof. In the working model, we propose that calvarial osteoblasts use WASL1-dependent lamellipodia to migrate toward the apex along a gradient of stiffness or adhesion of FN1 substrate (**Figure 6M**). Further, this mechanism is required for appropriate patterning of the calvaria and maintenance of the patency of the sutures between these bones.

## Discussion

The mechanisms by which the skull roof forms and is patterned in a determinative fashion is not understood. As these mechanisms underlie calvaria formation, it is attractive to ask if they also contribute to a common etiology of craniofacial disorders. Using a combination of genetic tools in the mouse, we develop a framework detailing the process of calvarial expansion and its impacts on suture patency. Central to these findings is that FN1 expression is graded towards the apex in the CM and is required for efficient apical expansion of the calvarial osteoblasts. Deletion of FN1 in the CM leads to altered cellular behaviors, without directly perturbing differentiation or proliferation of calvarial osteoblasts. Analysis of *Wasl* deficient mutants show that calvarial osteoblasts require lamellipodia-actin nucleation for integration of this graded substrate signal and apical expansion. Collectively, our data not only uncover the role FN1 plays in both growth of the calvaria and cranial suture maintenance, but also illuminate the possible relationship between calvarial growth and CS.

### Graded fibronectin as a substrate for frontal bone primordia growth

One hypothesis for directional growth of calvarial osteoblasts from the initial pool in the basal region of the SOM is the interpretation of graded environmental signals toward the apex that would provide directional cues. In support of this hypothesis, we observed graded FN1 expression between E11.5 to E13.5 along the baso-apical axis of the cranial mesenchyme (**Figure 1**). Mice with either elevated (Apert mutants) or loss (FN1 mutants) of FN1 in cranial mesenchyme have accelerated or diminished apical expansion of frontal bones, respectively. These findings suggest the graded FN1 expression may provide directional cues to guide the apical expansion of either individual or a collective group of calvaria osteoblasts in the frontal bone primordia. The absolute limits or steepness of that FN1 grade that is required for efficient apical expansion and timing of establishing the graded FN1 expression in the Apert mutants needs further investigation.

Directional cell movement along a graded matrix substrate can be guided by a combination of adhesion/haptotaxis, mechano/durotaxis, or chemotaxis as shown *in vitro* and *in vivo* (Shellard & Mayor, 2021a, 2021b). Fibroblasts and vascular smooth muscle cells migrate directionally *in vitro* in response to a mechanical stiffness gradient generated by fibronectin coated polyacrylamide hydrogel but not by laminin (Hartman et al., 2016; Lo et al., 2000; Phuong Le et al., 2022). Interesting, ARP2/3 is specifically required to direct differential dynamics of lamellipodia of cells moving up along a fibronectin gradient *in vitro* by haptotaxis (King et al., 2016). This functional dependency on ARP2/3 in interpretation of fibronectin structure synergizes with our finding that loss of *Wasl* function, which regulates APR2/3, leads to inability of calvarial osteoblasts to expand toward the apex. Future studies will focus on whether either or both these strategies are in play for apical expansion of calvaria and how the graded expression of FN1 is established in cranial mesenchyme using new models.

### Building a framework for directional cell movement during apical expansion of calvaria

Altogether, our data support a model in which calvarial osteoblasts expand apically by directional migration. There is no single model to explain the complex movement and regulation of directional migration in a proliferating primordium as multiple factors can be espoused as components of directional migration (Petrie et al., 2009). The variety of complex mesenchymal cell behaviors during migration within the rich 3D interstitial matrix *in vivo* are just emerging and still rely heavily on our understanding of *in vitro* studies. In order to build a conceptual framework for apical expansion of calvaria observed from *in vivo* conditional genetic mutants in this study, we can begin to draw connections from common themes that emerge from *in vitro* studies. First, there is consensus that mesenchymal cells migrate by using ARP2/3-branched actin nucleated by WASL at the leading edge to generate lamellipodia and adhere to a matrix substrate (Caswell & Zech, 2018; Doyle et al., 2009; Yamada & Sixt, 2019). Here, we found that filamentous actin is enriched and anisotropic in organization in the domain of frontal bone primordia, and *Wasl* is preferentially required in a cell autonomous manner for apical expansion of frontal bone osteoblasts. Second, there are features of the ECM matrix, such as graded expression, stiffness, and alignment that also shed light on how ECM proteins regulate directional cellular movement over long distances (Shellard & Mayor, 2021a; Yamada et al., 2019). We demonstrate that the FN1 extracellular matrix is graded in the cranial mesenchyme towards the apex and is required for the efficient apical expansion of frontal bones in the calvaria. Future studies will need to focus on the organization of FN1 fibrils and association with active cell movement in an *in vivo* context. Third, there are cellular behaviors indicative of mesenchymal cell movement that we show is directionally coordinated. Migrating mesenchymal cells and nuclei are elongated with their length-width ratio that is greater than two and with their actin cytoskeleton aligned to the ECM (Charras & Sahai, 2014; Faurobert et al., 2015; Friedl & Mayor, 2017; Petrie & Yamada, 2016; te Boekhorst et al., 2016; van Helvert et al., 2018). We found that the shapes of calvarial osteoblast nuclei become progressively more elongated along the baso-apical direction in the controls showing directional bias. This elongation is significantly reduced in the *Fn1* and *Wasl* mutants, as are the levels of associated F-actin fibers in the frontal bone primordia (**Figure 6M**). Fourth, many migrating cell types *in vitro* and in early development are polarized along the axis of growth which can be visualized by the asymmetric position of the Golgi complex at the motile end in front of the nuclei or in some contexts in the rearward position (Carney & Couve, 1989; Magdalena et al., 2003; Nemere et al., 1985; Serrador et al., 1999). Here, we found that the Golgi complex in controls was positioned forward and rearward of the nuclei of the primordia and the position became more biased forward of the nuclei in the apical region along the axis of growth. Further supporting directional growth in the formation of the cranium, the asymmetric bias of the Golgi complex in calvarial osteoblasts was significantly altered in the CM-*Fn1* mutants suggesting change in polarization after the loss of the FN1 substrate. Previous studies demonstrate that geometrical constraints of the FN1 matrix are key determinants of the Golgi complex position, cell polarity, and cell shape (Serrador et al., 1999). Altogether, our data from *in vivo* studies support a model where calvarial osteoblasts orient and elongate towards the apex and migrate using WASL-dependent lamellipodia on a graded FN1 substrate leading to apical expansion calvaria during development.

### Role of fibronectin in coronal suture maintenance

There are a large number of genes in which mutation can lead to CS in humans (Twigg & Wilkie, 2015). This disparate set of factors suggest that diverse mechanisms contribute to the pathogenesis of CS. Conversely, the broad genetic heterogeneity might suggest shared broad mechanisms that integrate multiple classes of signals.

*Fn1* conditional mutants show coronal craniosynostosis. We found approximately 80% penetrance for CS in the coronal suture of the CM-Fn1^*fl/fl*^ embryos. The coronal suture is preferentially lost in the FN1 mutant, similar to many well-studied CS models, such as *Twist1*^+/-^ and Apert mutants. In the CM-Fn1^*fl/fl*^ mutant, we found ectopic mineralized tissue in the medial region of the suture. How FN1 preserves the suture patency of this tissue at the cellular level is unclear. Teng et al. (2018) found that the frontal bones and parietal bones show increased diagonal growth in *Twist1*^+/-^ zebrafish, such that the coronal suture is vulnerable to be affected in these conditions. Holmes and Basilico (2012) also suggested that the suture mesenchyme retains its identity in coronal sutures and ectopically undergoes osteogenesis in Apert *Fgfr2*^*+/S252W*^ mice. Through the analysis of various CS models, the known cellular mechanisms underlying CS are also different, which include loss of osteogenic-non osteogenic boundary integrity (Merrill et al., 2006), and altered differentiation of suture mesenchyme in embryos (Holmes et al., 2009; Holmes & Basilico, 2012). It is unclear if FN1 has a direct role within sutures to maintain suture patency or if the effect is indirect. One hypothesis in line with these studies is that calvaria growth defects in both the Apert and conditional FN1 mutants lead to inappropriately timed suture formation and loss of signals to maintain patency.

The common presentation of suture fusion in genetically heterogeneous cases of CS suggests that shared developmental mechanisms may underlie disease pathology. Stamper et al. (2011) suggested that perturbations in the gene expression related to ECM-mediated focal adhesion occur in cases of non-syndromic single-suture craniosynostosis. Furthermore, during tissue development, ECM can directly guide the cell migration as a “road”, but ECM could also “stop” the initiation of the movement or hold the cells back at the destination (Walma & Yamada, 2020). By serving to direct migration, ECM can serve as a framework to unify the concentrations of cytokine or growth factors in lieu of a gradient during chemotaxis or haptotaxis (Colak-Champollion et al., 2019; Malhotra et al., 2018; Sieg et al., 2000), restrict cells as physical barriers (Renkawitz et al., 2019; Zanotelli et al., 2019), or modify cellular migration machinery through altering adhesion, protrusion or traction force generation (Richier et al., 2018; Sekine et al., 2012; Sieg et al., 2000; Walma & Yamada, 2020; Yamada & Sixt, 2019). Taken together our results demonstrate that FN1 can contribute to mammalian calvarial expansion and suture maintenance. These observations further provide a model in which the biological process governed by FN1 may serve as a convergence point downstream of other known pathways associated with CS disorders arriving at a similar phenotypic outcome. Our studies lend support to the concept that targeting extracellular matrix may be a novel avenue for treatment strategies for calvarial growth, fracture healing, and calvarial disorders.

## Acknowledgements

We thank all the members of the Atit Lab past and present for helpful comments and suggestions in guiding this project. We thank Nyoka Lovelace and Mia Carr for support with animal husbandry. This work was supported by NIH-NIDRCR: R56 DE030206 (RA, MH, GH) and R21: DE029348 (RA, SH). CWRU-SOURCE (MG), CWRU-ENGAGE (JL). Thank you to Dr. Nicole Crown for the generous use of the Leica confocal and to the CWRU bio[box] facility for use of microscopy and shared instrumentation facilities in the Biology Dept.

## Methods

### Mice strains and husbandry

*PDGFRαCreER* (Jax stock#018280) (Rivers et al., 2008), *Fn1* ^*flox/flox*^ (*Fn*^*fl/fl*^) (Jax stock #029624) (Sakai et al., 2001), *Wasl*^*fl/fl*^ (MGI:3760279) (Cotta-de-Almeida et al., 2007; Hawkins et al., 2021), *Wnt1Cre2* (Jax stock #022137) (Danielian et al., 1998), *EIIaCre* (Lakso et al., 1996), FGFR2^NeoS252W^ (Wang et al., 2005) were used to study the role of fibronectin in calvaria and suture development. For timed matings, *PDGFRαCreER/+; Fn*^*fl/fl*^ males were crossed with *Fn*^*fl/fl*^ females, *Wnt1Cre2/+; Wasl*^*fl/fl*^ males were crossed with *Wasl*^*fl/fl*^ overnight. All mice were genotyped as described previously and maintained on a mixed genetic background except the Apert line is inbred and maintained on C57BL/6J (Hawkins et al., 2021; Sakai et al., 2001; Wang et al., 2005). To yield the desired crosses, the mice were time mated overnight and checked for vaginal plugs in the morning. The vaginal plug day was assigned as embryonic day 0.5 (E0.5). The pregnant dams were orally gavaged with 25μg Tamoxifen/g body weight at E9.5 and 12.5μg Tamoxifen/g body weight at E10.5 to induce the CreER recombination. Tamoxifen was dissolved in corn oil and gavaged on designated day. Embryos were collected and processed as previously described (Atit et al., 2006). For each experiment, a minimum of four mutants with litter-matched controls from at least three litters were studied unless otherwise noted. Case Western Reserve University Institutional Animal Care and Use Committee approved all animal procedures in accordance with AVMA guidelines (Protocol 2013–0156, Animal Welfare Assurance No. A3145–01).

### Histological stains, Immunohistochemistry, and Immunofluorescence

E12.5-E15.5 tissue was collected and fixed in 4% paraformaldehyde (PFA) at 4°C for 35-50 min, respectively, sucrose dehydrated, embedded in O.C.T Compound (Tissue-Tek Sakura, Sakura Finetek USA, Torrance, CA, USA) and cryopreserved at -80°C. The embryos were cryosectioned at 14μm in the coronal or transverse plane. Immunofluorescence (IF) on cryosections was performed by drying slides at room temperature, washing in 1X PBS, and blocking in 5-10% donkey or goat serum. Sections were incubated with primary antibodies overnight at 4°C, washed in 1X PBS, incubated with species-specific secondary antibodies for 1 hour at room temperature, washed with DAPI (0.5 μg/mL), and then mounted with Fluoroshield (Sigma F6182). Following primary antibodies were used: Collagen hybridizing peptide (CHP) (Hwang et al., 2017; Jussila et al., 2021) (1: 250; 3Helix BIO60), rabbit anti-Fibronectin antibody (1: 250; Abcam ab2413; RRID:AB_2262874), rabbit anti-Fibulin 1 (1: 250; Abcam ab230994), rabbit anti-Collagen I (1: 250; Abcam ab270993; RRID:AB_2927551), Collagen III (Rockland Labs 1:100), rabbit anti-Laminin (1: 250; Millipore L9393), rabbit anti-Osx (1:2000 or 1:4000; Abcam ab209484; RRID:AB_2892207), rabbit anti-Caspase 3 (1: 250; Abcam ab13847; RRID:AB_443014), Goat anti-Runx 2 (1:100; R&D Systems AF2006-SP; RRID:AB_2184528) and mouse anti-GM130 (1:100; Thermo Fisher Scientific BDB610822; RRID:AB_398141). Appropriate species-specific secondary antibodies included Alexa 488 (1:500, Invitrogen A32790; RRID: AB_2762833) and Alexa 594 (1:800 Invitrogen A11012; RRID: AB_2534079). For IF staining against GM130, Vector M.O.M kit (BMK2202) was used before incubated with primary antibody.

To quantify cell proliferation index, mice were administered 250ug EdU in PBS/10g of body weight by intraperitoneal injection 30 min. prior to sacrifice. Embryos were then collected, cryopreserved, and sectioned as described above. EdU was detected using Click-iTEdU Alexa Fluor 594 Imaging kit as per manufacturer directions (Invitrogen C10639). The slides were then stained with 1: 2000 DAPI for nuclei and mounted with Fluoroshield (Sigma F6182). The percent of proliferation cells in the frontal bone primordia were quantified using ImageJ/Fiji (see below). Alizarin staining on cryosections was performed on sections after post-fixing in 4% PFA, washing in 1xPBS for 5 min, quickly rinsing with ddH_2_O before incubating with 2% Alizarin-S staining solution (Sigma Aldrich, TMS-008-C) for 15 seconds. The slides were washed with ddH_2_O and mounted with Vectashield mounting medium (H-1400-10). RGB Trichrome staining was performed on paraffin sections were stained as described previously (Electron Microscopy Sciences, 26357-02; Thermo Fisher Scientific, F99-10; Sigma-Aldrich, A3157) (Gaytan et al., 2020; Jussila et al., 2022). The paraffin sections were stained with Masson’s Trichrome for mature Collagen I or with Van-Gieson’s staining for elastin according to the standard protocols. Alkaline phosphatase staining was performed on cryosections after rinsing in PBS and NTMT (100mM Tris, pH 9.4, 100mM NaCl, and 50mM MgCl_2_) for 5 minutes. Embryos were stained with 20μl/mL NBT/BCIP substrate (Roche 11681451001) in the dark for 5-10 min at room temperature. Then the slides were washed with 1X PBS and mounted with aqueous mounting medium (H-1400-10). Slides were scanned using Hamamatsu Nanozoomer S60 by 20X brightfield scanning.

Phalloidin staining at E13.5 coronal sections was performed by incubating 1X Phalloidin working solution (1X PBS, 10% BSA and Phalloidin-iFluor™ 488) 1000x conjugate in DMSO (1:1000; Cayman Chemical Company 20549) for 45 minutes. After washing in 1X PBS twice, the sections were stained with DAPI and mounted with Fluoroshield (Sigma F6182). The images were captured by the Leica STELLARIS 5 confocal microscope with 63X oil immersion objective ((Leica; HC PL APO 63x/1.40 OIL CS2) using Application suite X.

The brightfield images for alkaline phosphatase staining, Alizarin Red, Masson’s Trichrome staining, RGB Trichrome staining and Van-Gieson’s staining as well as immunofluorescent staining images for CHP, collagen III, Laminin, Fibronectin, Fibulin 1, OSX, Caspase 3 and Runx 2, and EdU staining were captured at 10X and 20X using Olympus BX60 microscope (Olympus UPIanFI 4/0.13) with CellSens Entry Software. Confocal images were captured at 63X oil immersion objective on Leica STELLARIS 5 (Leica; HC PL APO 63x/1.40 OIL CS2) using Application suite X software. Images were processed in Adobe Photoshop and Fiji/ImageJ.

### Whole-mount skeletal preparation and Alizarin Red staining

Embryos at E16.5 were fixed in 95% ethanol and acetone at 4°C overnight before staining as previously described (Ferguson et al., 2018). Briefly, the embryos were pre-cleared in 1% potassium hydroxide (KOH) (Fisher Scientific, 1310–58-3) solution for 1 hour and stained in 0.005% alizarin red (Sigma, A5533) dissolved in 1% KOH overnight at 4°C. Then the embryos were placed in 50% glycerol (Sigma Aldrich, F6428-40ML): 1% KOH until clear. The embryos were kept in 100% glycerol for long-term storage. Images of controls and mutants were obtained at the same magnification on Lieca MZ16F stereoscope.

### MicroCT

MicroCT images were taken from fixed specimens maintained in 70% Ethanol using a Bruker Skyscan 1173, 240-degree scan with 0.2rotational step, X-ray source voltage 70 kV and current sett 80 mA. Exposure time was 1500 msec. Resolution of scan was 7 voxels. Images were processed using Amira software package, version 6.0 (FEI).

### Imaging

The brightfield images for histological stains were captured using the Olympus BX60 microscope with a digital camera (Olympus, DP70) and CellSens Entry Software (Olympus Corporation 2011; Version 1.5) with a 10X or 20X objective (Olympus UPIanFI 4/0.13). White balance was performed before the images were captured. The immunofluorescence images were captured on the Olympus BX60 wide-field microscope with Olympus DP70 camera or Leica STELLARIS 5 confocal microscope with a 63X oil immersion objective (Leica; HC PL APO 63x/1.40 OIL CS2) using Leica Application Suite X software (Leica; 4.1.1.232273). The exposure was held constant between controls and the experimental groups. Wholemount skeletal preparations were imaged with Leica MZ16F stereoscope and Leica DFC490 camera with Leica software. Images were processed using Adobe Photoshop and Fiji/ImageJ and page set in Adobe InDesign.

### Quantification and histomorphometrics

At least 3 sections per embryo were used for histomorphometrics. Quantifying the percent of cells expressing markers such as OSX, RUNX2 and EdU), data was measured as the ratio of marker expressing number of cell to total DAPI stained nuclei number in the region of interest (ROI) boxes in the frontal bone primordia. Corrected fluorescence intensity was calculated from immunofluorescence images with Fiji/ImageJ. For fluorescence intensity (such as Fibronectin1, phalloidin), images were transformed to 8-bit and average of mean gray values of three circles on non-fluorescence regions was calculated as the background fluorescence (McCloy et al., 2014). The corrected fluorescence per ROI was calculated by following formula: Corrected fluorescence=Integrated Density- (Area of ROI×Average of background fluorescence).

The apical length of alkaline phosphatase expressing frontal bone primordia was measured using NDP view2 (Hamamatsu, U12388-01) with free-hand line and then normalized to the length of cranial length from eye lid to apex.

For GM130 Golgi-complex to nuclei angle distribution, the orientation of the images was adjusted to the midline of the brain (90°). A vector from GM130 stained Golgi complex to the center of the nucleus was drawn and the angle of the vector and the x axis (0°) was measured by Fiji/ImageJ. In order to avoid oversampling, the total numbers of the measurements from each embryo and from the same regions were kept similar by using the random selection function in Excel and graphed to wind-rose plots in Excel.

For cellular nuclei shape analysis, the nuclei were stained with DAPI (1:2000) and the images were captured with 63X oil immersion objective by the confocal microscope (Leica STELLARIS 5). The measurements related to cell shape analysis of the nuclei in apical, intermediate and basal regions of interest in the OSX+ frontal bone primordia were determined by CellProfiler (CellProfiler 3.1.8) from 4 controls and 4 mutants, respectively. Max Feret diameter and Min Feret diameter were obtained by CellProfiler (CellProfiler 3.1.8). In addition to the measurements automatically assessed by the CellProfiler pipeline, the length-width aspect ratio was obtained by 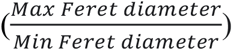. 25 nuclei per embryo for each measurement were randomly selected by Excel to avoid oversampling. Graphs and statistical tests were made by Prism 9.

Quantification of skeletal histomorphometries, the length and width of the E16.5 skulls were photographed in the dorsal view as normalization variables (labeled with red dash lines in Figure 6). The anterior and posterior metopic suture distances as well as the proportion of frontal bone area (colored in grey) to the frontal region (colored in pink) were measured. The graphs and statistics were generated by Prism 9.

For all images, the boundary of frontal bone primordia was defined on the expression of OSX, RUNX2 or alkaline phosphatase shown in the cranial mesenchyme region, or the morphological pattern discerned by the condensed DAPI staining and adjacent meningeal layers.

To quantify the alignment of the phalloidin stained F-actin filaments, single confocal microscope image of 0.5micron thickness of E12.5 frontal bone primordia were used. Three non-overlapping ROIs of fixed size in both the apical and basal regions of the frontal bone primordia in two histological sections of three biological replicates were analyzed. The ROIs were converted to 16-bit grayscale using FIJI/ImageJ and input into *Alignment by Fourier Transform (AFT)* (Marcotti et al., 2021), using MATLAB (Mathworks, v2022a). The following parameters were used for calculating an image order parameter: Window Size: 30 pixels, Window Overlap: 50%, Neighborhood Radius: 2x vectors. Local masking and filtering were not applied to the images. Vector angles distribution from an ROI can be visualized as a heat map and quantified collectively as an order parameter. Order parameter normally ranges between 0 for random alignment and 1 for perfectly aligned. Vector angles distribution from one ROI from each of the three embryos were combined and reported in the heat map. The median value of order parameter from three ROIs/section were averaged in the apical and basal region of two sections per embryo and graphed.

### RT-qPCR

The supraorbital cranial mesenchyme of E13.5 embryos was manually enriched and RNA was isolated as previously described (Ferguson et al., 2018; Hamburg-Shields et al., 2015). Relative mRNA expression was quantified using 4 ng of cDNA on a StepOne Plus Real-Time PCR System (Life Technologies) and the ΔΔCT method (Schmittgen & Livak, 2008). Commercially available TaqMan probes (Life Technologies) specific to each gene were used: *Fn1* (Mm01256744_m1) and β*-actin* (ActB, 4352933E). The CT values were normalized to β*-actin* CT value. ΔΔCT values were obtained by normalizing the ΔCT values to the average ΔCT values of the controls. Relative mRNA fold change was determined using the ΔΔCT values.

### Statistics

All graphs and statistical analysis were generated using Microsoft! Excel version 16.16.27 for Mac (Microsoft) and GraphPad Prism version 9.0.0 for Mac (GraphPad Software). Data are presented as mean ± SEM in all graphs. Outliers were excluded using the outlier calculator in Microsoft! Excel. The pairwise sample comparisons for cell proliferation, cell density and the quantification of FN1 and Phalloidin corrective fluorescence expression were performed using unpaired, two-tailed Welch’s t-test due to unequal variance and sample numbers. The results for GM130 analysis were compared using a Kolmogorov–Smirnov test developed for intracellular organization (Apte & Marshall, 2013). The p-values for statistical tests in all figures are represented as: *=p<0.05, **=p<0.01, ***=p<0.001 and ****=p<0.0001.

