## Supplementary figures for "Apical expansion of calvarial osteoblasts and suture patency is dependent on graded fibronectin cues"

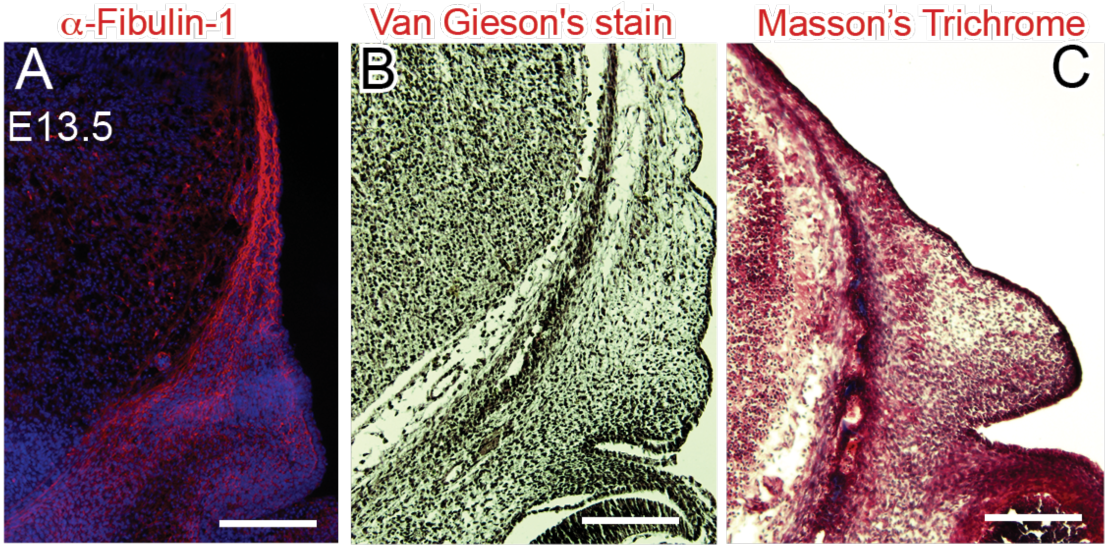


Supplementary Figure 1. (A) Immunofluorescence of Fibulin 1 in E13.5 coronal sections. In the fibrous network, Fibulin associates with assembled Fibronectin proteins. (B) Van Gieson's staining in E13.5 coronal sections showing the elastin is not enriched in cranial mesenchyme. (C) Masson's trichrome staining at E13.5 for mature collagen in cranial mesenchyme. Scale Bar= 100 μM.

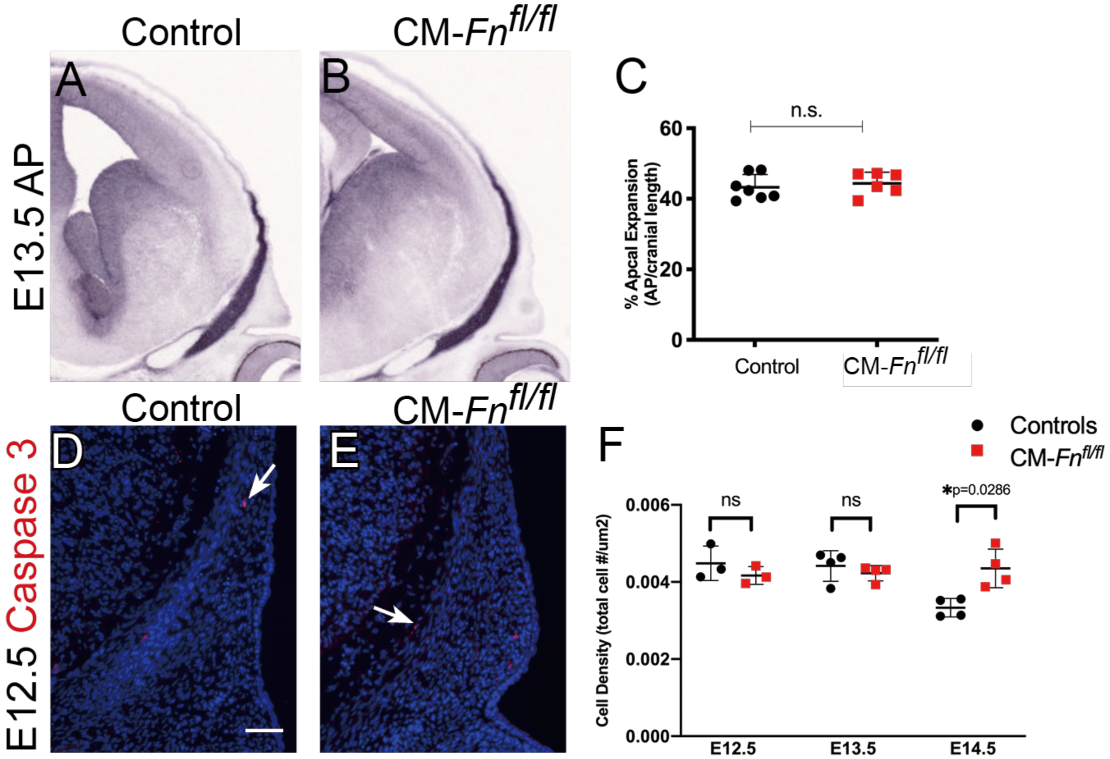


Supplementary Figure 3. **FN1 deletion does not affect cell density, cell death or apical expansion of the frontal bone primordia at E13.5.** (A, B, C) E13.5 CM-*Fn*$^{fl/fl}$ mutants show comparable apical expansion of frontal primordia to controls by Alkaline phosphatase staining. (D, E) Immunofluorescence of apoptosis marker activated Caspase 3 showing there is no change in cell death of E12.5 CM-*Fn*$^{fl/fl}$ mutants relative to controls. (F) The cell density of calvarial osteoblasts is increased at E14.5 in CM-*Fn*$^{fl/fl}$ mutants relative to controls whereas it is not perturbed at E12.5 or E13.5. N = 3-7 controls; 4-6 mutants. Scale Bar = 100 μM. Panel A and B are obtained from Nanozoomer at the same magnification.

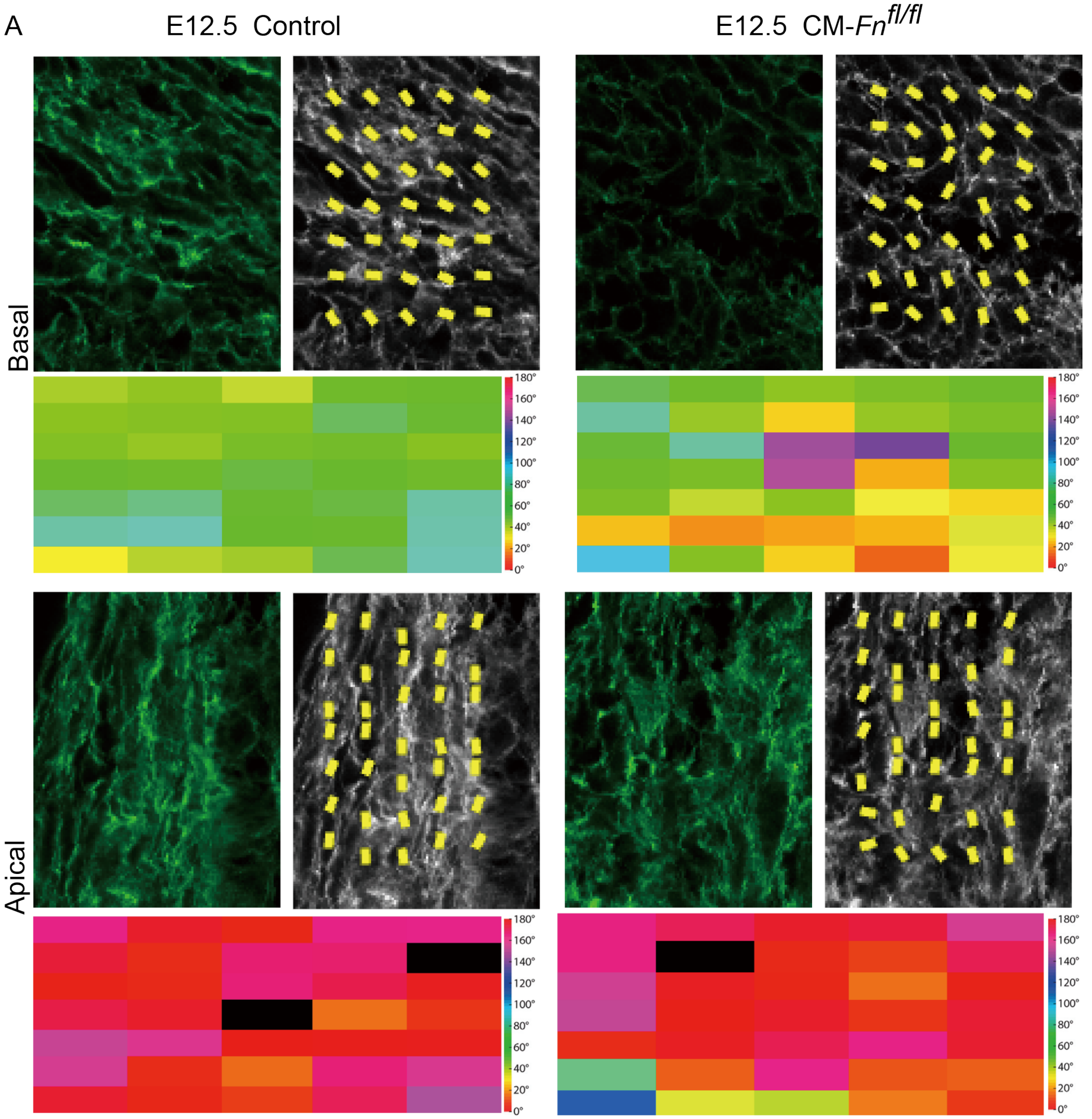


Supplementary Figure 4. **Calvarial osteoblasts in CM-*Fn*$^{fl/fl}$ shows altered alignment of F-actin in the basal region of frontal bone primordia at E12.5**. (A) The staining of phalloidin and corresponding heat maps and vector maps showing the orientation of F-actin fibers in apical and basal regions at E12.5, respectively. N = 3 controls, 3 mutants. The data were obtained from a ROI in the apical and basal regions of the frontal bone primordia of a control and a CM-*Fn*$^{fl/fl}$ mutant, respectively. All the images are obtained at the same magnification.

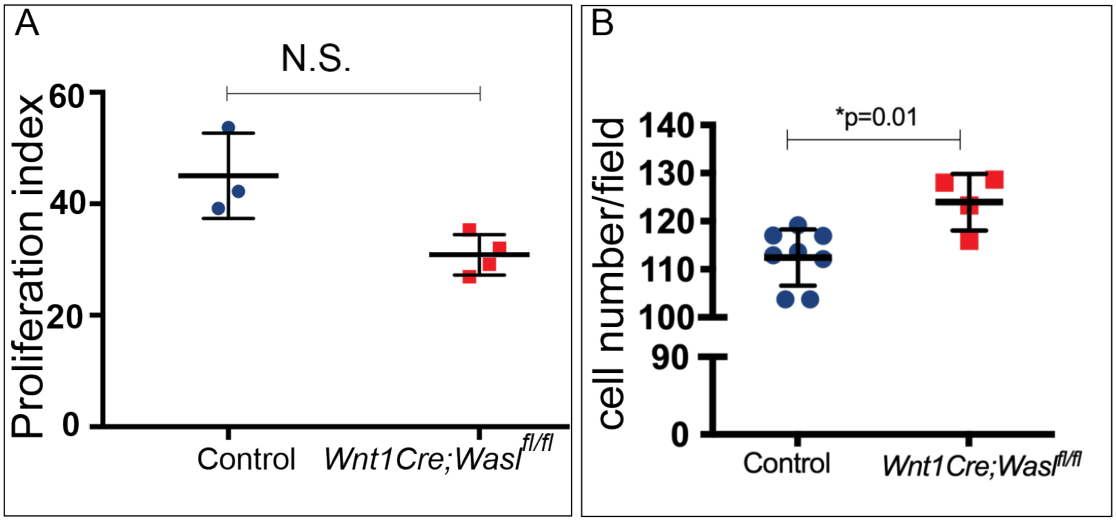


Supplementary Figure 5. ***Wnt1Cre; Wasl$^{fl/fl}$* mutants show comparable cell proliferation index and increased cell number in fixed field in the frontal bone primordia.** (A) The overall cell proliferation index is not altered in *Wnt1Cre; Wasl$^{fl/fl}$* mutants at E14.5, relative to the controls. N = 3 controls; 4 mutants. (B) *Wnt1Cre; Wasl$^{fl/fl}$* mutants show higher cell number in a fixed area in frontal bone primordia, relative to the controls at E14.5. N = 6 controls; 4 mutants.
